# Loci for prediction of penicillin and tetracycline susceptibility in *Neisseria gonorrhoeae*: a genome wide association study

**DOI:** 10.1101/2021.08.03.454909

**Authors:** Tatum D. Mortimer, Jessica J. Zhang, Kevin C. Ma, Yonatan H. Grad

**Author notes:** These authors contributed equally (Tatum D. Mortimer, Jessica J. Zhang). Corresponding author: Yonatan H. Grad, Department of Immunology and Infectious Diseases, Harvard T.H. Chan School of Public Health, 665 Huntington Ave, Building 1, Room 715, Boston, Massachusetts 02115.

## Abstract

**Background:** While *Neisseria gonorrhoeae* poses an urgent public health threat because of increasing antimicrobial resistance, much of the circulating population remains susceptible to historical treatment regimens. Point-of-care diagnostics that report susceptibility could allow for reintroduction of these regimens, but development of such diagnostics has been limited to ciprofloxacin, for which susceptibility can be predicted from a single locus.

**Methods:** We assembled a dataset of 12,045 *N. gonorrhoeae* genomes with phenotypic resistance data for tetracycline (n = 3,611) and penicillin (n = 6,935). Using conditional genome wide association studies (GWAS), we sought to define genetic variants associated with susceptibility to penicillin and tetracycline. We evaluated the sensitivity and specificity of these variants for predicting susceptibility and non-resistance in our collection of gonococcal genomes.

**Findings:** In our conditional penicillin GWAS, the presence of a genetic variant defined by a non-mosaic *penA* allele without an insertion at codon 345 was significantly associated with penicillin susceptibility and had the highest negative effect size of significant variants (*p* = 5.0 × 10^−14^, *β* = -2.5). In combination with the absence of *bla*_TEM_, this variant predicted penicillin susceptibility with high specificity (99.8%) and modest sensitivity (36.7%). For tetracycline, the wild type allele at *rpsJ* codon 57, encoding valine, was significantly associated with tetracycline susceptibility (*p* = 5.6 × 10^−16^, *β* = -1.6) after conditioning on the presence of *tetM*. The combination of *rpsJ* codon 57 allele and *tetM absence* predicted tetracycline susceptibility with high specificity (97.2%) and sensitivity (88.7%).

**Interpretation:** As few as two genetic loci can predict susceptibility to penicillin and tetracycline in *N. gonorrhoeae* with high specificity. Molecular point-of-care diagnostics targeting these loci have the potential to increase available treatments for gonorrhea.

**Funding:** National Institute of Allergy and Infectious Diseases, the National Science Foundation, and the Smith Family Foundation

**Research in Context:** *Evidence before this study:* We searched PubMed with the terms “*Neisseria gonorrhoeae*” and “diagnostic” or “assay” plus “penicillin” or “tetracycline” for reports in any language published up to July 1, 2021. We additionally searched for “*Neisseria gonorrhoeae*” and “genome wide association study”. We found that previously proposed molecular diagnostics for penicillin and tetracycline susceptibility either exclusively focused on plasmid-mediated resistance (i.e., targeting *bla*_TEM_ or *tetM*) or did not include variants in genes encoding antibiotic targets (e.g., did not include *penA* or *rpsJ*). Targets for molecular surveillance have focused on resistance-associated alleles rather than susceptibility-associated alleles. We did not find any previous penicillin or tetracycline conditional genome wide association studies (GWAS) in *N. gonorrhoeae*.

*Added value of this study:* To identify targets for molecular diagnostics that predict penicillin and tetracycline susceptibility, we conducted GWAS conditioning on the presence of plasmid-mediated resistance determinants to detect chromosomal loci with the highest association with susceptibility. We discovered a sequence (*penA*_01) that differentiates susceptible isolates from those with a resistance-associated insertion at codon 345 and from those with mosaic *penA* alleles, which is associated with penicillin susceptibility. We also found that *rpsJ* codon 57 was the chromosomal locus contributing the most to tetracycline susceptibility. The combination of these chromosomal loci and the absence of plasmid encoded determinants predicts penicillin and tetracycline susceptibility with high specificity in both a large global collection of *N. gonorrhoeae* and a validation dataset consisting of recently published genomes from CDC’s Gonococcal Isolate Surveillance Program (GISP) surveillance collected in 2018.

*Implications of all the available evidence:* The chromosomal loci *penA_*01 and *rpsJ* codon 57 in combination with plasmid loci *bla*_TEM_ and *tetM* are candidates for the development of point-of-care molecular diagnostics for penicillin and tetracycline susceptibility. The loci may be combined with the currently available ciprofloxacin susceptibility diagnostics to predict susceptibility to multiple antibiotics. Additionally, our study suggests that conditional GWAS focused on variants associated with susceptibility may be a promising approach to identify minimal sets of loci for molecular diagnostics and surveillance.

## Introduction

Gonorrhea, caused by infection with *Neisseria gonorrhoeae*, is the second most reported notifiable infection in the United States at a rate of 188.4 cases per 100,000 people in 2019 and increasing antibiotic resistance has made it an urgent public health threat.^1^ Treatment is empiric, and resistance has limited the recommended treatment in the US to ceftriaxone, an extended spectrum cephalosporin (ESC).^2^

Despite the emergence of multidrug resistant strains,^3^ a large fraction of clinical isolates remain susceptible to multiple antibiotics.^1^ Data from the Gonococcal Isolate Surveillance Project (GISP), the US Centers for Disease Control and Prevention’s sentinel surveillance system for antibiotic resistance in *N. gonorrhoeae*, reported that in 2019 44.5% of clinical isolates were not resistant to any tested antibiotics, 64.6% were non-resistant (defined as minimum inhibitory concentrations (MIC) corresponding to susceptible or intermediate categories), to ciprofloxacin (MIC < 1 µg/mL), 72.2% were non-resistant to tetracycline (MIC < 2 µg/mL), and 87.2% were non-resistant to penicillin (MIC < 2 µg/mL).^1^

Point-of-care diagnostics that inform on antibiotic susceptibility may help forestall the emergence and spread of resistance by enabling a shift from empiric to tailored treatment and expanding the number of antibiotics used to treat *N. gonorrhoeae* infections.^4^ The observation that ciprofloxacin susceptibility can be predicted with high specificity and sensitivity based on *gyrA* codon 91 has led to the development of molecular tests that query this locus; the SpeeDx ResistancePlus GC, for example, was recently approved for clinical use in Europe and granted ‘breakthrough designation’ by the FDA.^5^ However, extension of this sequence-based approach to other antibiotics has been stymied, as they lack single locus determinants of susceptibility and resistance.

Penicillin (PCN) and tetracycline (TET) were the recommended therapies for gonorrhea until the 1980s, when the prevalence of high level resistance increased enough to prompt a switch in the empiric treatment regimen.^6,7^ Resistance to PCN and TET can be both chromosomal and plasmid mediated. Chromosomally-encoded resistance arises from mutations modifying the antibiotic targets—*rpsJ*^8^ for TET resistance and *penA*^9,10^ and *ponA*^11^ for PCN—and mutations in the porin *porB* and in the efflux pump *mtr* operon^12^. The plasmid-borne beta lactamase *bla*_TEM_^13^ confers high level PCN resistance and the ribosome protection protein *tetM*^14^ confers TET resistance. Despite previously being first-line gonorrhea treatments for decades, molecular diagnostics for PCN and TET susceptibility have been less commonly studied. Proposed diagnostics or targets of molecular surveillance for PCN susceptibility have focused on *bla*_TEM,_^15^ which performs poorly in the setting of chromosomally-encoded resistance, *porB*,^16^ which neglects important target modifying mutations in *penA*, or resistance-associated *penA* alleles^17^ rather than susceptibility-associated alleles. Similarly, assays targeting *tetM* have been developed, but they have not incorporated chromosomally-encoded tetracycline resistance.^15^

While there are multiple pathways to resistance for each drug, the key goal for sequence-based diagnostics is to predict susceptibility—rather than resistance—with high specificity. Therefore, here we sought to identify a concise set of loci that are associated with PCN and TET susceptibility using genome wide association studies (GWAS) and evaluate their predictive performance in gonococcal clinical isolates.

## Methods

### Study design and datasets

We collected publicly available whole genome sequencing (WGS) data (n = 12,045) and PCN (n = 6,935) and TET MICs (n = 5,727) from clinical *N. gonorrhoeae* isolates. For 2,116 isolates, TET MICs were reported as ≤ 4 µg/mL or ≤ 8 µg/mL. These MICs were excluded from further analyses, since we could not classify them as susceptible or resistant. To validate our results, we assembled 1479 genomes from CDC’s 2018 Gonococcal Isolate Surveillance Program (GISP) collection,^18^ representing the first five viable isolates collected each month from urethral specimens at sentinel surveillance sites across the United States. This study used publicly available data and did not require IRB approval.

### Procedures

Pipelines for genome assembly (Supplementary Figure 1, appendix p. 5) and resistance-associated allele (Supplementary Table 1, appendix p. 8) calling are detailed in the Appendix and follow previously described methods.^19^

### Statistical analysis

To identify variants associated with PCN and TET susceptibility, we performed conditional GWAS^20^ incorporating the presence of high effect size plasmid-mediated resistance (appendix pp. 2-5). The GWAS employed a linear mixed model and were run using Pyseer v 1.2.0^21^ with default allele frequency filters using unitigs (unique sequences representing SNPs, insertions, deletions, and changes in gene content) as genetic variants^22^. We also repeated the GWAS with kmers as genetic variants to ensure that the unitig calling procedure did not impact our results. Most datasets reported PCN MICs within the range of 0.06 - 32 μg/mL. Isolates with PCN MICs reported as ‘>4’ or ‘>2’ were not included in the GWAS analysis since the precise MIC was unknown; the final PCN GWAS dataset size was 6220 isolates after excluding isolates with missing genotypic or phenotypic data. Similarly isolates with imprecise TET MICs were excluded (e.g. “≤ 4”, “≤ 8”); the final dataset size for the TET GWAS was 3,453 isolates after excluding isolates with missing genotypic or phenotypic data. The GWAS incorporated isolate dataset of origin, country of origin, and presence of plasmid-encoded resistance determinants (*bla*_TEM,_ *tetM*) as fixed effect covariates. A similarity matrix was included as a random effect to correct for population structure. The significance of variants was assessed using a likelihood ratio test. We additionally corrected for multiple hypothesis testing using a Bonferroni correction based on the number of unique presence/absence patterns for unitigs or kmers. The threshold for significance in the penicillin GWAS was 3.13 × 10^−7^ for unitigs and 3.49 × 10^−8^ for kmers, and the threshold for significance in the tetracycline GWAS was 3.41 × 10^−7^ for unitigs and 4.44 × 10^−8^ for kmers.

We evaluated the sensitivity and specificity of susceptibility-associated alleles to predict PCN and TET susceptibility using CLSI breakpoints for susceptibility (PCN MIC ≤ 0.06 µg/mL, TET MIC ≤ 0.25 µg/mL) and non-resistance (susceptible or intermediate, PCN MIC < 2 µg/mL, TET MIC < 2 µg/mL) in both the global and validation datasets. We additionally used isolate metadata from the 2018 GISP collection to estimate the prevalence of isolates with susceptibility associated genotypes across patient groups (sexual behavior, race/ethnicity). χ^2^ tests were performed in R v 4.0.3^23^ using infer v 0.5.4 (https://infer.tidymodels.org/) using a threshold for significance of p < 0.05. Confidence intervals for sensitivity and specificity were calculated using the formula 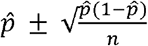, where 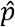 is sensitivity or specificity and *n*is the number of true positives or true negatives, respectively.^24^

## Results

Since plasmid-encoded resistance determinants are known to contribute to high level resistance for both PCN and TET, we used conditional GWAS to identify additional variants contributing to susceptibility. Here, we focused on significant variants associated with increased susceptibility (i.e., negative effect size, *β*). We found that a unitig (*penA*_01, Supplementary Figure 2A, appendix p. 6) corresponding to non-mosaic *penA* alleles without the resistance-associated insertion at codon 345 was significantly associated with PCN susceptibility (Supplementary Figure 3A, appendix p. 7, *p* = 5.0 × 10^−14^, *β* = -2.5). After conditioning on the presence of *tetM*, we found that a unitig (Supplementary Figure 2B, appendix p. 6) corresponding to the wild-type allele at *rpsJ* codon 57, encoding valine, was significantly associated with TET susceptibility (Supplementary Figure 3B, appendix p. 7, *p* = 5.6 × 10^−16^, *β* = -1.6). Significant unitigs also mapped to *porB* (PCN: *p =* 2.0 × 10^−23^, *β =* - 0.60; TET: *p =* 2.5 × 10^−50^, *β =* -0.49) and a loss of function variant in *mtrC* (PCN: *p =* 2.5 × 10^−50^, *β =* -1.2; TET: *p =* 1.1 × 10^−14^, *β =* -1.0) for both antibiotics; however, effect sizes (*β*) were lower than unitigs mapping to antibiotic targets. We found that using kmers as the genetic variant instead of unitigs did not impact the results. The significant kmers with the largest effect on penicillin susceptibility (*p* = 5.3 × 10^−14^, *β* = -2.5) overlapped the *penA_*01 unitig, and the significant kmers with the largest effect on tetracycline susceptibility (*p* = 4.4 × 10^−16^, *β* = -1.6) overlapped the wild-type *rpsJ* 57 unitig.

We used the presence of *penA*_01 combined with the absence of *bla*_TEM_ to predict PCN susceptibility in our global dataset (Figure 1A). We found that this susceptibility-associated genotype predicted PCN susceptibility and non-resistance with high specificity but low sensitivity (Table 1). For TET susceptibility prediction, we identified isolates with the wild-type allele at *rpsJ* codon 57 combined with the absence of *tetM* (Figure 1B). This combination predicted TET susceptibility and non-resistance with high specificity and sensitivity (Table 1). The addition of one chromosomal marker improves performance, as prediction of susceptibility based on plasmid-encoded determinants alone had low sensitivity in our dataset (Supplementary Table 2, appendix p. 8).

**Figure 1.**
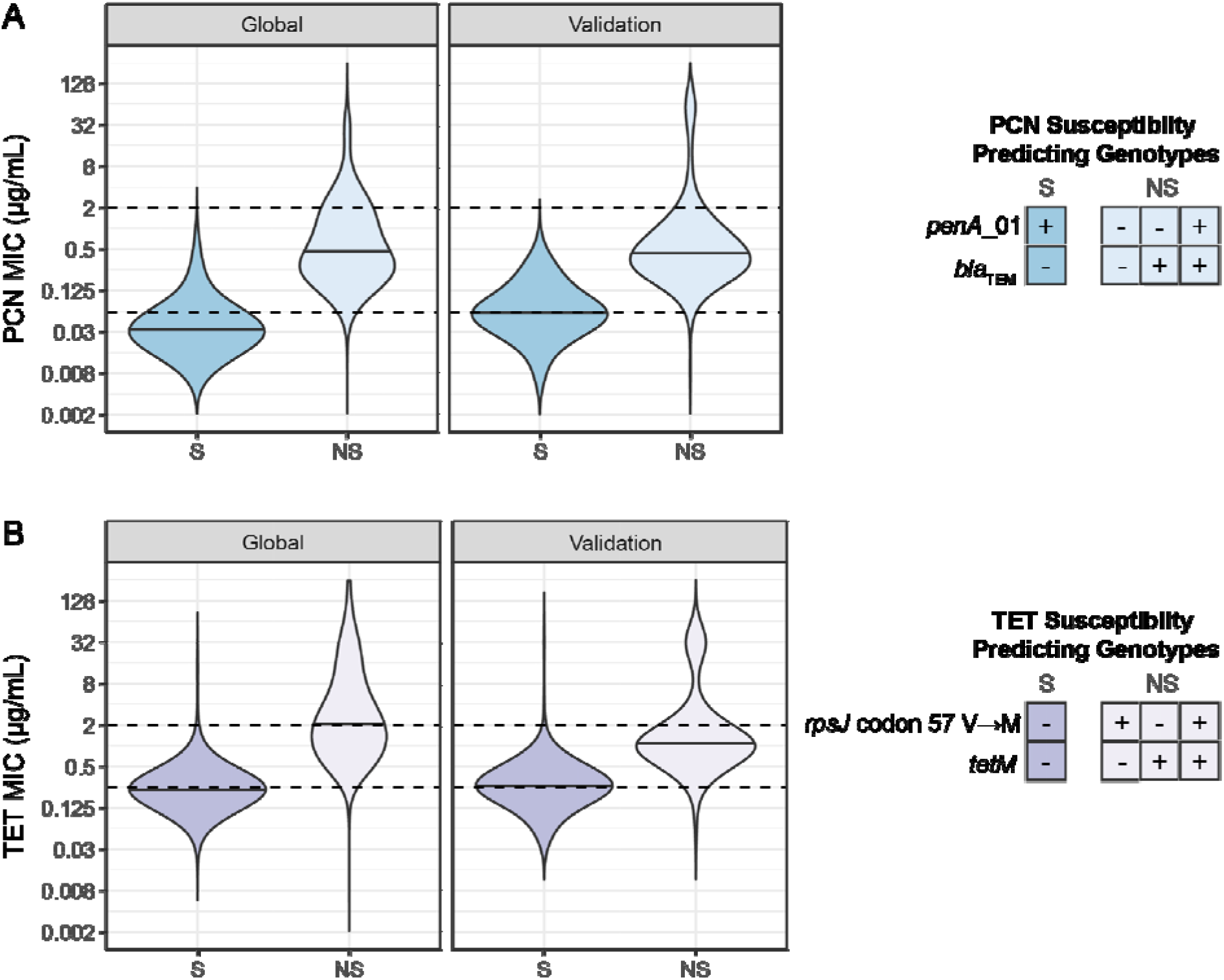
Penicillin and tetracycline MICs are lower in isolates with susceptibility associated genotypes in global and validation datasets. Dashed lines indicate CLSI breakpoints for susceptibility and resistance. A) Penicillin MICs of isolates with *penA*_01 and without *blaTEM* (S, dark blue) compared to isolates with one or more of these determinants (NS, light blue). For the global dataset, MICs from 6,935 isolates are plotted; 111 had the susceptible (S) genotype and 6824 had the non-susceptible (NS) genotype. For the validation dataset, MICs from 1,479 are plotted; 57 had the S genotype, and 1422 had the NS genotype. B) Tetracycline MICs of isolates with wild-type *rpsJ* (57V) and without *tetM* (S, dark purple) compared to isolates with one or more of these determinants (NS, light purple). For the global dataset, MICs from 3,611 isolates are plotted; 497 had the susceptible (S) genotype and 3114 had the non-susceptible (NS) genotype. For the validation dataset, MICs from 1,477 are plotted; 245 had the S genotype, and 1,232 had the NS genotype.

**Table 1.** Sensitivity and specificity of genotypes for predicting PCN and TET susceptibility, with 95% confidence interval (CI).

| Genotype | Phenotype | Global Dataset |  | Validation Dataset (GISP 2018<sup>18</sup>) |  |
| --- | --- | --- | --- | --- | --- |
|  |  | Sensitivity (95% CI) | Specificity (95% CI) | Sensitivity (95% CI) | Specificity (95% CI) |
| *penA*_01 + absence of *bla*<sub>TEM</sub> | PCN susceptible (MIC ≤ 0.06 µg/mL) | 36.7%<br>(31.8% - 41.5%) | 99.8%<br>(99.4% - 100.0%) | 63.6%<br>(56.2% - 71.1%) | 98.9%<br>(97.4% - 100.0%) |
|  | PCN non-resistant (MIC < 2 µg/mL) | 2.1%<br>(0.7% - 3.4%) | 100.0%<br>(100.0% - 100.0%) | 4.4%<br>(1.7% - 7.1%) | 100.0%<br>(100.0% - 100.0%) |
| *rpsJ* WT + absence of *tetM* | TET susceptible (MIC ≤ 0.25 µg/mL) | 88.7%<br>(87.2% - 90.3%) | 97.2%<br>(96.4% - 98.0%) | 78.2%<br>(75.1% - 81.3%) | 94.9%<br>(93.2% - 96.5%) |
|  | TET non-resistant (MIC < 2 µg/mL) | 28.3%<br>(26.3% - 30.3%) | 99.7%<br>(99.4% - 99.9%) | 22.1%<br>(19.4% - 24.8%) | 99.5%<br>(99.0% - 99.9%) |

Since PCN and TET MICs were not reported for all isolates, we identified these mutations in our full genomic dataset: 2.1% (252/12,045) had the PCN susceptibility-associated genotype, and 15.9% (1951/12,045) of isolates had the TET susceptibility-associated genotype. The prevalence of these genotypes varied across genomic epidemiology studies (Supplementary Table 3, appendix p. 9). The majority of isolates with non-susceptible genotypes encode only chromosomal resistance determinants. Among isolates with PCN non-susceptible genotypes, 14.7% (1,734/11,793) encoded *bla*_TEM_. 19.3% (1636/8491) of isolates with TET non-susceptible genotypes encoded *tetM*.

To validate our observations in a relatively unbiased dataset from the United States, we assembled a recently published collection of *N. gonorrhoeae* genomes from CDC’s Gonococcal Isolate Surveillance Program^18^; in this collection, isolates were not selected for sequencing based on their susceptibility phenotypes. First, we verified that the *penA* sequence identified in the GWAS (*penA_*01) also identified isolates with non-mosaic *penA* alleles without the 345 insertion in the validation dataset. In this dataset, 100% (57/57) of isolates with the *penA_*01 encoded non-mosaic *penA* alleles without the insertion when the full length *penA* allele was examined.

We also calculated sensitivity and specificity for prediction of PCN and TET susceptibility and non-resistance in the GISP collection (Figure 1, Table 1). In two isolates, we were unable to genotype *rpsJ* codon 57 because of insufficient coverage of either the reference or alternate allele. Similar to results from the global collection, specificity was high for both antibiotics and CLSI cutoffs. Sensitivity increased for PCN prediction and decreased for TET prediction, reflecting different proportions of isolates with MICs at the CLSI breakpoints in the global and validation datasets compared to the number of true positives in the dataset. For example, 88.3% (151/171) of false negative isolates in the global dataset have MICs at the breakpoint of 0.06 µg/mL, and the global dataset contains a lower proportion of susceptible isolates, with only 99 true positives (Supplementary Table 4, appendix p. 10).

In addition to antimicrobial resistance phenotypes, GISP reports information on patient characteristics for each isolate collected. To analyze the utility of these genotypic markers in different patient populations, we calculated the prevalence of the susceptibility-associated genotypes across patient groups. Susceptible genotypes were more common among men who have sex with women (MSW) compared to men who have sex with men (MSM) and men who have sex with men and women (MSMW) for both PCN (χ^2^ test, df = 3, p = 0.0035) and TET (χ^2^ test, df = 3, p < 0.0001). The prevalence of the PCN susceptibility-associated genotype was 5.2% (44/853), 1.5% (7/479), and 2.2% (2/91), in MSW, MSM, and MSMW, respectively. For TET, the susceptibility-associated genotype was 20.6% in MSW (175/851), 9.6% in MSM (46/479), and 9.9% (9/91) in MSMW. Additionally, the susceptibility-associated genotypes varied across race and ethnicity groups and were enriched in samples from Black men; however, prevalence of susceptibility-associated genotypes were not significantly different between race and ethnicity groups when MSM and MSW were considered separately (Supplementary Table 5, appendix p. 11).

## Discussion

Here, we have used conditional GWAS incorporating known, high effect size variants^20^ to identify targets for diagnostics addressing both plasmid and chromosomally mediated PCN and TET resistance. We found that the combination of *penA*_01, representing non-mosaic *penA*^9^ without an insertion at codon 345^10^, and the absence of *bla*_TEM_ predicts PCN susceptibility and that the combination of *rpsJ* codon 57^8^ and the absence of *tetM* predicts TET susceptibility. These loci defined the most susceptible isolates in our dataset and predicted susceptibility (PCN MIC ≤ 0.06 µg/mL, TET MIC ≤ 0.25 µg/mL) with high specificity to both antibiotics in both our global dataset and an unbiased collection from the United States. Sensitivity was high for tetracycline susceptibility prediction and modest for penicillin susceptibility prediction.

Given that many gonorrhea infections are diagnosed by molecular tests and culture and subsequent MIC testing require multiple days, gonorrhea infections are currently treated empirically based on population levels of resistance. Point-of-care diagnostics are a potential approach for targeted therapy of gonorrhea in the future. The results here suggest that, of the many possible chromosomal loci to predict penicillin and tetracycline susceptibility, *penA*_01 and *rpsJ* are promising targets for diagnostic development. As currently available molecular diagnostics, including SpeeDx ResistancePlus GC^5^ and Xpert MTB/RIF^25^, target multiple loci, we expect that a diagnostic incorporating the loci identified here in addition to *gyrA* 91 (comprising five total loci) could be developed using existing technology to provide susceptibility information for three antibiotics. These loci could additionally be used for culture-independent molecular epidemiology and surveillance, as WGS directly from patient samples is not currently routine. Typing schemes such as NG-STAR^26^ targeting resistance determinants have been developed; however, these schemes have not focused on loci specific to penicillin and tetracycline resistance.

Utility of a diagnostic or molecular surveillance targeting these loci may vary in different patient populations. For example, the prevalence of susceptibility associated genotypes varied across genomic epidemiology studies included in our global dataset, reflecting both enrichment of antibiotic resistant isolates in some studies as well as variable selection pressure from antibiotic use in different regions. WGS data from *N. gonorrhoeae* isolated in the United States, Europe, and Australia make up the majority of available genomic data, and the composition of the *N. gonorrhoeae* population in other regions is unknown. Similar to other studies of the association between *N. gonorrhoeae* antibiotic resistance and patient demographics, prevalence of these susceptibility-associated genotypes vary across patient groups defined by sexual behavior and race/ethnicity in isolates collected by GISP.^27–29^ In the United States, a diagnostic for PCN and TET susceptibility may be most useful in populations with higher prevalence of infection with susceptible isolates, such as MSW and women.

Beyond the uneven sampling mentioned above, our study has additional limitations. While we assign isolates as susceptible based on MIC, MIC measurements may vary by up to two doubling dilutions, which makes the categorization of isolates with MICs near the breakpoint potentially more prone to error; however, errors of this magnitude are rare.^30^ We focused on identifying a single chromosomal locus to combine with the absence of plasmid-encoded determinants and predict susceptibility. The addition of other loci (e.g., *mtr* and *porB*) may be needed to increase sensitivity for the higher cutoff (MIC < 2 µg/mL) but with as yet unclear impact on specificity.

The alleles identified here from genomic analyses are promising targets for the development of POC molecular diagnostics for *N. gonorrhoeae* susceptibility to penicillin and tetracycline. Diagnostics that evaluate as few as two loci per drug could allow for the reintroduction into clinical use of these gonococcal treatment regimens. The impact of test sensitivity on treatment options and prevalence of antibiotic resistance as well as the impact of querying additional loci are important avenues for future research and further development of sequence-based diagnostics of antimicrobial susceptibility.

## Author Contributions

TDM and JJZ performed the GWAS and statistical analyses. TDM and KCM assembled genomic dataset. YHG supervised and managed the study. All authors contributed to data interpretation. TDM, JJZ, and YHG wrote the manuscript. All authors reviewed and edited the final manuscript. TDM and JJZ had full access to the data in the study, and all authors were responsible for the decision to submit for publication.

## Acknowledgments

TDM is supported by NIH NIAID (F32AI145157). YHG is supported by R01 AI132606 and R01 AI153521 and by the Smith Family Foundation. KCM is supported by NSF GRFP grant DGE1745303.

## Data Sharing Statement

The analysis pipeline and data are available at github.com/gradlab/pcn_tet_susceptibility_gwas

