## appendix for "Loci for prediction of penicillin and tetracycline susceptibility in *Neisseria gonorrhoeae*: a genome wide association study"

### Table of Contents

Supplementary Methods. 2

Genome assembly. 2

Identification of known resistance associated alleles. 2

Genome wide association studies. 2

Model. 2

Phenotype. 3

Unitig and kmer calling. 3

Population structure. 3

Covariates. 3

Significance testing. 3

Annotation of significant variants. 3

Supplementary Figures. 5

Supplementary Figure 1. Mapping and assembly statistics. 5

Supplementary Figure 2. Significant susceptibility-associated unitigs aligned to representative resistance-associated sequences. 6

Supplementary Figure 3. Conditional GWAS identifies target-site variants associated with susceptibility to PCN and TET. 7

Supplementary Tables. 8

Supplementary Table 1. Previously described penicillin and tetracycline resistance markers. 8

Supplementary Table 2. Sensitivity and specificity of plasmid-associated markers for predicting PCN and TET susceptibility with 95% confidence interval. 8

Supplementary Table 3. Prevalence of susceptibility associated genotypes across datasets. 9

Supplementary Table 4. Counts for true positives, false positives, true negatives, and false negatives in the global and validation datasets. 10

Supplementary Table 5. Number and percentage of GISP isolates reported from 2018 encoding susceptibility associated genotypes by sexual behavior and race/ethnicity. 11

References. 12

### Supplementary Methods.

##### Genome assembly.

Sequencing reads from publicly available *N. gonorrhoeae* datasets (Supplementary Table 2) were downloaded from the European Nucleotide Archive. Quality of reads was assessed using FastQC (https://www.bioinformatics.babraham.ac.uk/projects/fastqc/). Reads were mapped to the NCCP11945 (NC_011035.1) reference genome using BWA-MEM v 0.7.17[1]. Duplicate reads were marked with Picard v 2.20.1 (https://broadinstitute.github.io/picard/), and reads were sorted with samtools v X. The quality of mapped reads was assessed using Qualimap's bamqc v 2.2.1[2]. We used Pilon v 1.23[3] to call variants (minimum mapping quality 20 and minimum coverage of 10). To create pseudogenomes, we replaced the reference allele with high quality variant calls (at least 90% of reads supporting allele). Positions called as deletions by Pilon were replace with a gap character ('-'), and positions with low coverage or an indeterminant allele were replaced with an N.

Sequencing reads were additionally assembled using SPAdes v 3.12.0[4]. Assemblies were corrected by mapping reads to contigs using the --careful option. Contigs less than 500bp long or with less than 10X coverage were removed from assemblies. Quality of assemblies was assessed using QUAST v 5.0.2 [5] (Supplementary Figure 3B).

Accessions and quality control statistics are available at github.com/gradlab/pcn_tet_susceptibility_gwas in the data/prediction/ directory for the global dataset and data/validation for the GISP 2018[6] dataset.

##### Identification of known resistance associated alleles.

Alleles previously described to be associated with resistance (Supplementary Table 4) were identified in pseudogenomes or *de novo* assemblies. Single nucleotide variants occurring in the background of *N. gonorrhoeae* alleles were called using Pilon variant calls (e.g. genomic position 2031479 for *rpsJ* codon 57). *penA* alleles from assemblies were typed according to the naming scheme and mosaic designations used by the NG-STAR database (last accessed March 2, 2021)[7], requiring 100% identity to be matched to a specific allele. The resistance associated insertion in *penA* at codon 345 was identified using blastn from BLAST+ v 2.9.0[8] of *de novo* assemblies with alleles from FA1090 (NC_002946.2) serving at the wild type reference. The presence of plasmid-mediated resistance elements ($bla_{\text{TEM}}$ and *tetM*) were also identified using blastn of *de novo* assemblies; the query sequences were NG_068038.1 and MG874353.1 accession, respectively.

The presence of resistance alleles are available at github.com/gradlab/pcn_tet_susceptibility_gwas in the data/prediction/ directory for the global dataset and data/validation for the GISP 2018[6] dataset.

##### Genome wide association studies.

As in previous studies[9,10], we used a linear mixed model based GWAS implemented in pyseer v 1.3.6[11] to test for associations of variants with penicillin and tetracycline MICs.

*Model.*

$$Y \sim \mathrm{W}\alpha + \mathrm{X}\beta + \mathrm{u} + \epsilon$$

$$u \sim N\left(0, \sigma_g^2 K\right)$$

$$\epsilon \sim N(0, \sigma_e^2 I)$$

$Y$ is a vector of MICs. W and α are the covariate matrix and their fixed effects, respectively. X is a vector representing the presence of the unitig or kmer, and β is the fixed effect for the genetic variant. To control for population structure, $u$ is a random effect parameterized with $K$ (a similarity matrix) and $\sigma_g^2$ (additive genetic variance). Non-genetic effects are modeled by the random effect $\epsilon$.

*Phenotype.*

For 2116 isolates with available MICs, TET MICs were reported as ≤ 4 μg/mL or ≤ 8 μg/mL. These were excluded since we could not classify them as susceptible or resistant. Isolates with PCN MICs reported as '>4' or '>2' were not included in the GWAS analysis since the precise MIC was unknown. 6220 isolates were included in the penicillin GWAS, and 3453 isolates were included in the tetracycline GWAS. For all other datasets, when reported MICs contained '>' or '<' symbols, these symbols were stripped and the numerical value was used as the MIC for GWAS. MICs were log2-transformed.

*Unitig and kmer calling.*
Unitigs and kmers were generated from de novo assemblies to represent the genetic variation present within this dataset. Unitigs were generated from a compressed de Bruijn graph with unitig-counter v 1.1.0 (https://github.com/johnlees/unitig-counter), based on the approach used in DBGWAS[12]. Kmers were generated using fsm-lite v 1.0 (https://github.com/nvalimak/fsm-lite). Unitigs and kmers present in 1%-99% of the dataset were tested for association with the phenotype.

*Population structure.*
We used an alignment of pseudogenomes for phylogenetic analysis. To identify recombinant regions and estimate a recombination free phylogeny, we used Gubbins v 2.4.1[13] and RAxML v 8.2.12[14]. A similarity matrix describing the relatedness of isolates was generated flag from the phylogeny using phylogeny_distance.py from pyseer with the --lmm flag.

*Covariates.*
We included the country of isolation and original dataset as covariates in the GWAS to correct for geographic differences in MIC protocols and study specific sequencing artifacts. We also included the presence of plasmid-mediated resistance determinants encoded as 0 for absent and 1 for present as covariates.

*Significance testing.*
The significance of variants was assessed using a likelihood ratio test. We additionally corrected for multiple hypothesis testing using a Bonferroni correction based on the number of unique presence/absence patterns for unitigs or kmers obtained from count_patterns.py from pyseer. The threshold for significance in the penicillin GWAS was $3.13 \times 10^{-7}$ (0.05/159609) for unitigs and $3.49 \times 10^{-8}$ (0.05/1433207) for kmers, and the threshold for significance in the tetracycline GWAS was $3.41 \times 10^{-7}$ (0.05/146496) for unitigs and $4.44 \times 10^{-8}$ (0.05/1125008) for kmers.

*Annotation of significant variants.*
Unitigs and kmers were mapped to the WHO_N reference genome (GCA_900087725.2) as this genome contains both the *tetM*-encoding conjugative plasmid and the $bla_{TEM}$-encoding plasmid with pyseer's annotate_hits_pyseer. The output was used to generate Manhattan plots in R v 4.0.3[15] with ggplot2[16]. We considered significant unitigs and kmers that uniquely mapped to one location in the genome for further analyses.

To visualize variation between susceptibility associated unitigs and the homologous region in resistant isolates, we aligned *penA*_01, which is present in WHO_F (NZ_LT591897.1) to the equivalent *penA* region in isolates encoding the insertion at codon 345 (WHO_N, NZ_LT591910.1), mosaic *penA* 10

(WHO_K, NZ_LT591908.1 ), mosaic *penA* 34 (WHO_Y, NZ_LT592161.1), and mosaic *penA* 60 (FC428, NZ_AP018377.1). We also aligned the tetracycline susceptibility associated unitig, which is present in WHO_F to the equivalent *rpsJ* region in WHO_N. These alignments were visualized with Jalview 2.11.1.4[17].

### Supplementary Figures.

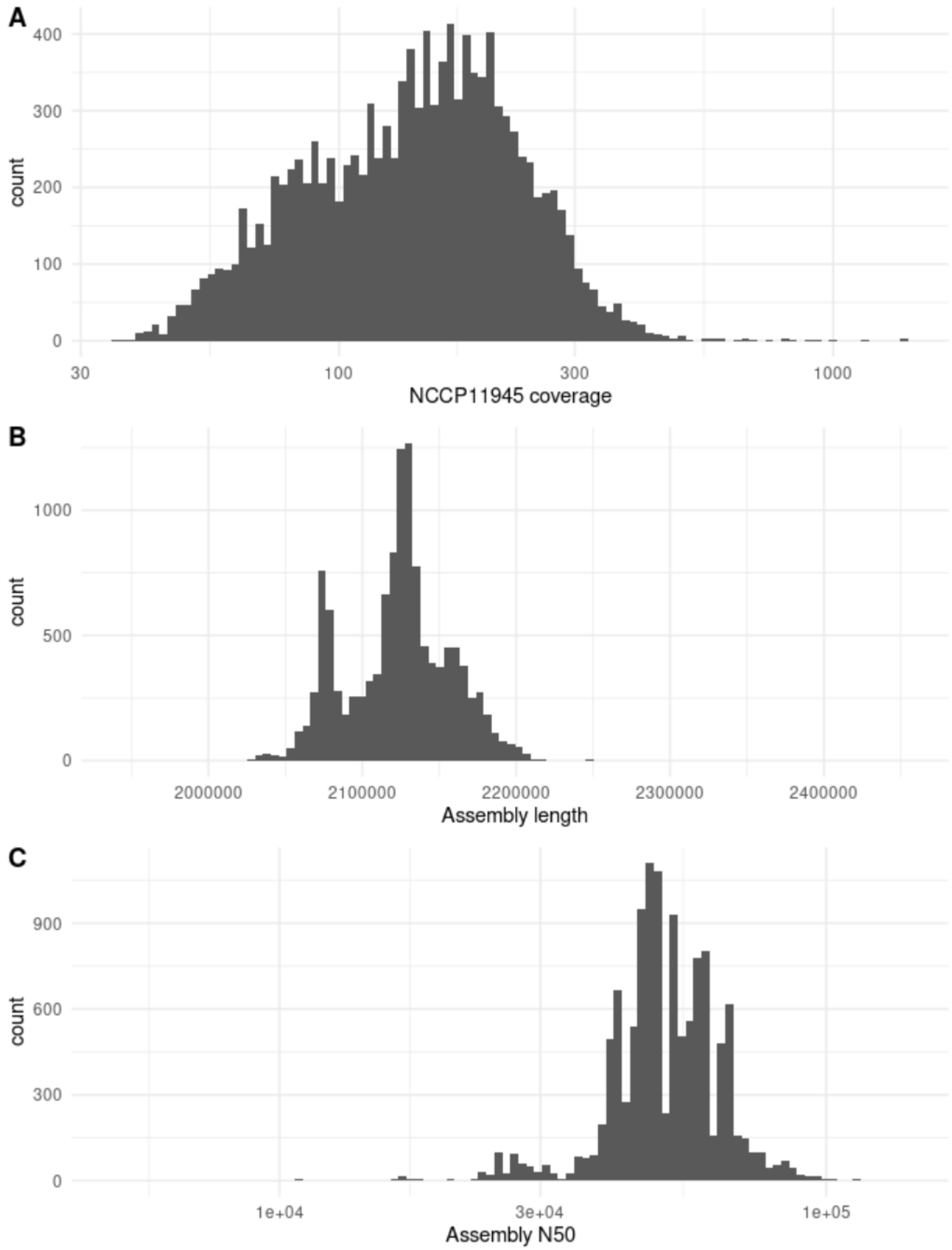


**Supplementary Figure 1. Mapping and assembly statistics.** Quality control was performed with Qualimap v and Quast v for *de novo* assemblies of sequencing data from 12,045 *N. gonorrhoeae* isolates. **A)** Coverage of NCCP11945 reference genome after mapping with bwa-mem **B)** Total assembly length after filtering contigs <500bp or <10X coverage **C)** Assembly N50

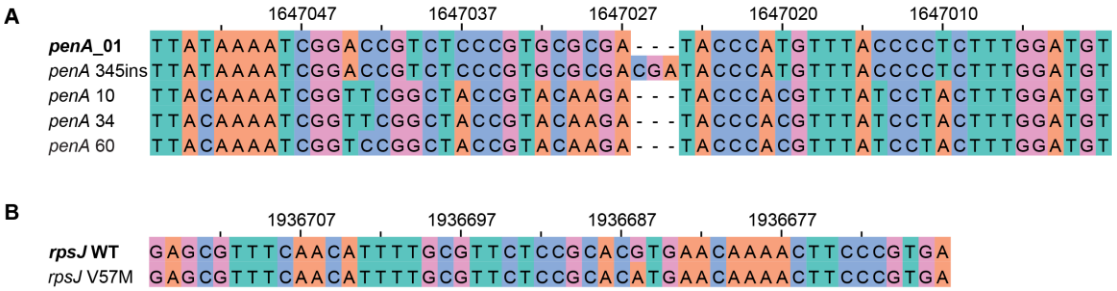


**Supplementary Figure 2. Significant susceptibility-associated unitigs aligned to representative resistance-associated sequences.** Indexing is relative to the WHO_F reference sequence (RefSeq: NZ_LT591897.1). Both *penA* and *rpsJ* are encoded on the negative strand in WHO_F. A) Penicillin susceptibility associated *penA*_01 (WHO_F 1647000 - 1647056) aligned to a non-mosaic *penA* with a resistance-associated insertion at codon 345 (WHO_N 1459748 - 1459807) and three common mosaic *penA* alleles, *penA* 10 (WHO_K 1462123 - 1462179), *penA* 34 (WHO_Y 1566575 - 1566631), and *penA* 60 (FC428 1556944 - 1557000). B) Tetracycline susceptibility *rpsJ* WT unitig (WHO_F 1936667 - 1936716) aligned to *rpsJ* with G to A nucleotide variant corresponding to V57M (WHO_N 1971964 - 1972013).

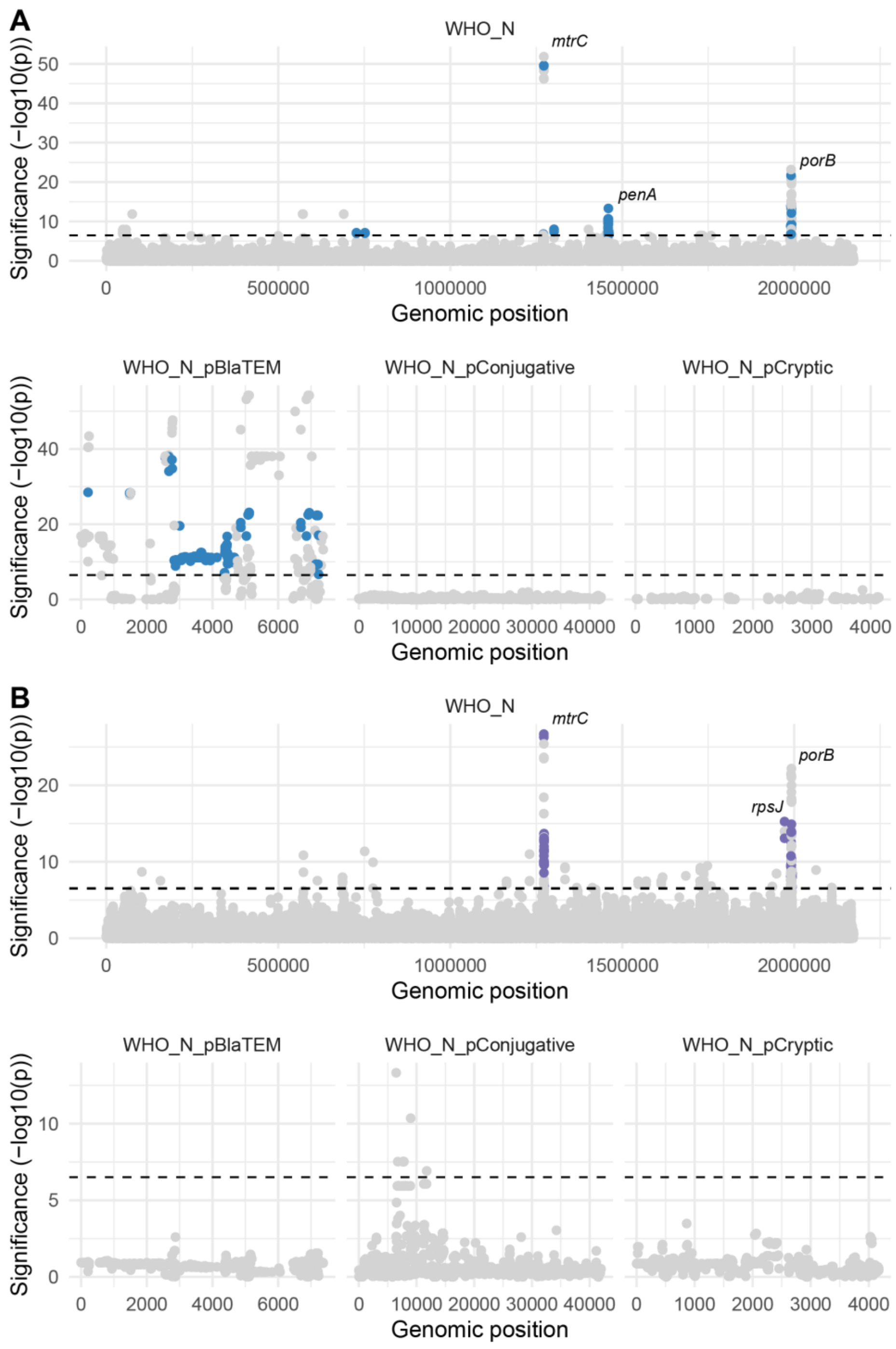


**Supplementary Figure 3. Conditional GWAS identifies target-site variants associated with susceptibility to PCN and TET.** Manhattan plots show unitig significance (-$\log_{10}$ transformed *p*-values) and unitig location determined by mapping to the WHO N reference genome, including the chromosome, $bla_{\text{TEM}}$ encoding plasmid, *tetM* encoding conjugative plasmid, and cryptic plasmid. The significance threshold is denoted by the horizontal, dashed line. **A)** Manhattan plot of unitig (n = 124,070) association with PCN susceptibility. Unitigs significantly associated with increased PCN susceptibility ($\beta < 0$) are blue. **B)** Manhattan plot of unitig (n = 158,087) association with TET susceptibility. Unitigs significantly associated with increased TET susceptibility ($\beta < 0$) are purple.

### Supplementary Tables.

#### Supplementary Table 1. Previously described penicillin and tetracycline resistance markers.

| Antibiotic | Gene | Variants |
|---|---|---|
| Penicillin | *bla*$_{TEM}$[41] | |
| | *ponA* | L421P[42] |
| | *penA* | Mosaic alleles[43]<br>345ins[44,45]<br>A501V/P[46]<br>G545S/I312M/V316T[47] |
| Tetracycline | *tetM*[48] | |
| | *rpsJ* | V57M[49] |
| Both | *porB* | G120K, A121D/N[50,51] |
| | *mtr* operon | promoter[52]<br>*mtrR* A39T, G45D[53] |

#### Supplementary Table 2. Sensitivity and specificity of plasmid-associated markers for predicting PCN and TET susceptibility with 95% confidence interval.

| Genotype | Phenotype | Sensitivity (95% CI) | Specificity (95% CI) |
|---|---|---|---|
| Absence of *bla*$_{TEM}$ | PCN susceptible (MIC ≤ 0.06 μg/mL) | 98.5% (97.8% - 99.3%) | 13.9% (11.8% - 16.0%) |
| | PCN non-resistant (MIC < 2 μg/mL) | 98.3% (98.1% - 98.5%) | 51.8% (51.1% - 52.5%) |
| Absence of *tetM* | TET susceptible (MIC ≤ 0.25 μg/mL) | 99.1% (98.7% - 99.5%) | 23.4% (21.4% - 25.4%) |
| | TET non-resistant (MIC < 2 μg/mL) | 99.4% (99.1% - 99.6%) | 40.0% (37.8% - 40.2%) |

**Supplementary Table 3. Prevalence of susceptibility associated genotypes across datasets.**

The susceptible genotype for PCN is the presence of penA_01 and the absence of blaTEM. The susceptible genotype for TET is the presence of wild-type rpsJ codon 57 and the absence of tetM. Non-susceptible genotypes include all other genotypes.

| Dataset | PCN genotypes | | TET genotypes | |
|---|---|---|---|---|
| | Susceptible | Non-susceptible | Susceptible | Non-susceptible |
| Alfsnes et al. 2020 [18] | 18 (1.9%) | 906 (98.1%) | 222 (24.0%) | 702 (76.0%) |
| Buckley et al. 2018 [19] | 0 (0.0%) | 93 (100.0%) | 93 (100.0%) | 0 (0.0%) |
| Cehovin et al. 2018 [20] | 0 (0.0%) | 103 (100.0%) | 0 (0.0%) | 103 (100.0%) |
| De Silva et al. 2016 [21] | 35 (2.5%) | 1365 (97.5%) | 219 (15.6%) | 1181 (84.4%) |
| Demczuk et al. 2015 [22] | 7 (6.6%) | 99 (93.4%) | 20 (18.9%) | 86 (81.1%) |
| Demczuk et al. 2016 [23] | 3 (1.5%) | 193 (98.5%) | 1 (0.5%) | 195 (99.5%) |
| Eyre et al. 2017 [24] | 21 (9.1%) | 210 (90.9%) | 33 (14.3%) | 198 (85.7%) |
| Ezewudo et al. 2015 [25] | 3 (5.6%) | 51 (94.4%) | 8 (14.8%) | 46 (85.2%) |
| Fifer et al. 2018 [26] | 0 (0.0%) | 50 (100.0%) | 0 (0.0%) | 50 (100.0%) |
| Grad et al. 2014, 2016 [27,28] | 11 (1.0%) | 1084 (99.0%) | 56 (5.1%) | 1039 (94.9%) |
| Harris et al. 2018 [29] | 54 (5.3%) | 960 (94.7%) | 119 (11.7%) | 895 (88.3%) |
| Kwong et al. 2017 [30] | 0 (0.0%) | 94 (100.0%) | 2 (2.1%) | 92 (97.9%) |
| Lan et al. 2020 [31] | 2 (0.9%) | 226 (99.1%) | 2 (0.9%) | 226 (99.1%) |
| Lee et al. 2018 [32] | 4 (1.0%) | 393 (99.0%) | 119 (30.0%) | 278 (70.0%) |
| Mortimer et al. 2020 [33] | 34 (3.8%) | 862 (96.2%) | 129 (14.4%) | 767 (85.6%) |
| Peng et al. 2019 [34] | 0 (0.0%) | 421 (100.0%) | 0 (0.0%) | 421 (100.0%) |
| Ryan et al. 2018 [35] | 0 (0.0%) | 39 (100.0%) | 1 (2.6%) | 38 (97.4%) |
| Sánchez-Busó et al. 2018 [36] | 30 (7.9%) | 348 (92.1%) | 36 (9.5%) | 342 (90.5%) |
| Thomas et al. 2019 [37] | 7 (1.1%) | 606 (98.9%) | 59 (9.6%) | 554 (90.4%) |
| Town et al. 2020 [38] | 13 (1.0%) | 1259 (99.0%) | 257 (20.2%) | 1016 (79.8%) |
| Williamson et al. 2019 [39] | 6 (0.3%) | 2175 (99.7%) | 506 (23.2%) | 1675 (76.8%) |
| Yahara et al. 2018 [40] | 4 (1.5%) | 256 (98.5%) | 36 (13.8%) | 224 (86.2%) |

**Supplementary Table 4. Counts for true positives, false positives, true negatives, and false negatives in the global and validation datasets.**

| Genotype | Phenotype | Global Dataset | | | | Validation Dataset (GISP 2018[6]) | | | |
|---|---|---|---|---|---|---|---|---|---|
| | | True Positives | False Positives | True Negatives | False Negatives | True Positives | False Positives | True Negatives | False Negatives |
| *penA*_01 + absence of *bla*$_{TEM}$ | PCN susceptible (MIC ≤ 0.06 µg/mL) | 99 | 12 | 6653 | 171 | 42 | 15 | 1398 | 24 |
| | PCN non-resistant (MIC < 2 µg/mL) | 111 | 0 | 1617 | 5207 | 57 | 0 | 190 | 1232 |
| *rpsJ* WT + absence of *tetM* | TET susceptible (MIC ≤ 0.25 µg/mL) | 409 | 88 | 3062 | 52 | 183 | 62 | 1181 | 51 |
| | TET non-resistant (MIC < 2 µg/mL) | 491 | 6 | 1869 | 1245 | 243 | 2 | 378 | 854 |

**Supplementary Table 5. Number and percentage of GISP isolates reported from 2018 encoding susceptibility associated genotypes by sexual behavior and race/ethnicity.** Dash indicates that no isolates were collected from the demographic group indicated. The susceptible genotype for PCN is the presence of *penA*_01 and the absence of $bla_{TEM}$. The susceptible genotype for TET is the presence of wild-type *rpsJ* codon 57 and the absence of *tetM*.

| | **PCN Susceptible Genotype** | | | **TET Susceptible Genotype** | | |
|---|---|---|---|---|---|---|
| | **MSM** | **MSMW** | **MSW** | **MSM** | **MSMW** | **MSW** |
| American Indian | 0 (0.0%) | - | 0 (0.0%) | 0 (0.0%) | - | 2 (40.0%) |
| Asian | 0 (0.0%) | 2 (5.1%) | 1 (5.0%) | 1 (5.3%) | 0 (0.0%) | 5 (25.0%) |
| Black | 20 (5.0%) | 0 (0.0%) | 33 (5.7%) | 13 (10.8%) | 3 (7.7%) | 32 (22.9%) |
| Hispanic | 1 (0.9%) | 0 (0.0%) | 1 (0.9%) | 13 (11.8%) | 1 (7.7%) | 13 (11.6%) |
| Multi-racial | 0 (0.0%) | 0 (0.0%) | 2 (10.0%) | 1 (6.25%) | 0 (0.0%) | 4 (20.0%) |
| Native Hawaiian | 0 (0.0%) | 0 (0.0%) | 0 (0.0%) | 1 (11.1%) | 0 (0.0%) | 0 (0.0%) |
| White | 0 (0.0%) | 0 (0.0%) | 7 (6.8%) | 14 (8.0%) | 3 (11.5%) | 17 (16.5%) |
| p-value ($\chi^2$ test) | 0.073 | | 0.96 | 0.52 | | 0.19 |

### References.

1 Li H. Aligning sequence reads, clone sequences and assembly contigs with BWA-MEM. *arXiv:13033997 [q-bio]* 2013; published online March 16. http://arxiv.org/abs/1303.3997 (accessed Sept 29, 2015).

2 Okonechnikov K, Conesa A, García-Alcalde F. Qualimap 2: advanced multi-sample quality control for high-throughput sequencing data. *Bioinformatics* 2015; : btv566.

3 Walker BJ, Abeel T, Shea T, *et al.* Pilon: An Integrated Tool for Comprehensive Microbial Variant Detection and Genome Assembly Improvement. *PLOS ONE* 2014; **9**: e112963.

4 Bankevich A, Nurk S, Antipov D, *et al.* SPAdes: A New Genome Assembly Algorithm and Its Applications to Single-Cell Sequencing. *Journal of Computational Biology* 2012; **19**: 455–77.

5 Gurevich A, Saveliev V, Vyahhi N, Tesler G. QUAST: quality assessment tool for genome assemblies. *Bioinformatics* 2013; **29**: 1072–5.

6 Reimche JL, Chivukula VL, Schmerer MW, *et al.* Genomic analysis of the predominant strains and antimicrobial resistance determinants within 1479 Neisseria gonorrhoeae isolates from the U.S. Gonococcal Isolate Surveillance Project in 2018. *Sex Transm Dis* 2021; published online May 14. DOI:10.1097/OLQ.0000000000001471.

7 Demczuk W, Sidhu S, Unemo M, *et al.* Neisseria gonorrhoeae Sequence Typing for Antimicrobial Resistance, a Novel Antimicrobial Resistance Multilocus Typing Scheme for Tracking Global Dissemination of N. gonorrhoeae Strains. *J Clin Microbiol* 2017; **55**: 1454–68.

8 Camacho C, Coulouris G, Avagyan V, *et al.* BLAST+: architecture and applications. *BMC Bioinformatics* 2009; **10**: 421.

9 Ma KC, Mortimer TD, Hicks AL, *et al.* Adaptation to the cervical environment is associated with increased antibiotic susceptibility in Neisseria gonorrhoeae. *Nature Communications* 2020; **11**: 4126.

10 Ma KC, Mortimer TD, Duckett MA, *et al.* Increased power from conditional bacterial genome-wide association identifies macrolide resistance mutations in Neisseria gonorrhoeae. *Nature Communications* 2020; **11**: 5374.

11 Lees JA, Galardini M, Bentley SD, Weiser JN, Corander J. pyseer: a comprehensive tool for microbial pangenome-wide association studies. *Bioinformatics* 2018; **34**: 4310–2.

12 Jaillard M, Lima L, Tournoud M, *et al.* A fast and agnostic method for bacterial genome-wide association studies: Bridging the gap between k-mers and genetic events. *PLOS Genetics* 2018; **14**: e1007758.

13 Croucher NJ, Page AJ, Connor TR, *et al.* Rapid phylogenetic analysis of large samples of recombinant bacterial whole genome sequences using Gubbins. *Nucl Acids Res* 2015; **43**: e15–e15.

14 Stamatakis A. RAxML version 8: a tool for phylogenetic analysis and post-analysis of large phylogenies. *Bioinformatics* 2014; **30**: 1312–3.

15 R Core Team. R: A Language and Environment for Statistical Computing. Vienna, Austria: R Foundation for Statistical Computing, 2021 https://www.R-project.org/.

16 Wickham H. ggplot2: Elegant Graphics for Data Analysis. Springer-Verlag New York, 2016 https://ggplot2.tidyverse.org.

17 Waterhouse AM, Procter JB, Martin DMA, Clamp M, Barton GJ. Jalview Version 2—a multiple sequence alignment editor and analysis workbench. *Bioinformatics* 2009; **25**: 1189–91.

18 Alfsnes K, Eldholm V, Olsen AO, *et al.* Genomic epidemiology and population structure of Neisseria gonorrhoeae in Norway, 2016–2017. *Microbial Genomics* 2020. DOI:10.1099/mgen.0.000359.

19 Buckley C, Forde BM, Trembizki E, Lahra MM, Beatson SA, Whiley DM. Use of whole genome sequencing to investigate an increase in Neisseria gonorrhoeae infection among women in urban areas of Australia. *Scientific Reports* 2018; **8**: 1503.

20 Cehovin A, Harrison OB, Lewis SB, *et al.* Identification of Novel Neisseria gonorrhoeae Lineages Harboring Resistance Plasmids in Coastal Kenya. *J Infect Dis* 2018; **218**: 801–8.

21 De Silva D, Peters J, Cole K, *et al.* Whole-genome sequencing to determine transmission of Neisseria gonorrhoeae: an observational study. *The Lancet Infectious Diseases* 2016; **16**: 1295–303.

22 Demczuk W, Lynch T, Martin I, *et al.* Whole-Genome Phylogenomic Heterogeneity of Neisseria gonorrhoeae Isolates with Decreased Cephalosporin Susceptibility Collected in Canada between 1989 and 2013. *J Clin Microbiol* 2015; **53**: 191–200.

23 Demczuk W, Martin I, Peterson S, *et al.* Genomic Epidemiology and Molecular Resistance Mechanisms of Azithromycin-Resistant Neisseria gonorrhoeae in Canada from 1997 to 2014. *J Clin Microbiol* 2016; **54**: 1304–13.

24 Eyre DW, De Silva D, Cole K, *et al.* WGS to predict antibiotic MICs for Neisseria gonorrhoeae. *J Antimicrob Chemother* 2017; **72**: 1937–47.

25 Ezewudo MN, Joseph SJ, Castillo-Ramirez S, *et al.* Population structure of Neisseria gonorrhoeae based on whole genome data and its relationship with antibiotic resistance. *PeerJ* 2015; **3**: e806.

26 Fifer H, Cole M, Hughes G, *et al.* Sustained transmission of high-level azithromycin-resistant Neisseria gonorrhoeae in England: an observational study. *Lancet Infect Dis* 2018; **18**: 573–81.

27 Grad YH, Kirkcaldy RD, Trees D, *et al.* Genomic epidemiology of Neisseria gonorrhoeae with reduced susceptibility to cefixime in the USA: a retrospective observational study. *The Lancet Infectious Diseases* 2014; **14**: 220–6.

28 Grad YH, Harris SR, Kirkcaldy RD, *et al.* Genomic Epidemiology of Gonococcal Resistance to Extended-Spectrum Cephalosporins, Macrolides, and Fluoroquinolones in the United States, 2000–2013. *J Infect Dis* 2016; **214**: 1579–87.

29 Harris SR, Cole MJ, Spiteri G, *et al.* Public health surveillance of multidrug-resistant clones of Neisseria gonorrhoeae in Europe: a genomic survey. *The Lancet Infectious Diseases* 2018; **18**: 758–68.

30 Kwong JC, Chow EPF, Stevens K, *et al.* Whole-genome sequencing reveals transmission of gonococcal antibiotic resistance among men who have sex with men: an observational study. *Sex Transm Infect* 2017; : sextrans-2017-053287.

31 Lan PT, Golparian D, Ringlander J, Van Hung L, Van Thuong N, Unemo M. Genomic analysis and antimicrobial resistance of Neisseria gonorrhoeae isolates from Vietnam in 2011 and 2015–16. *J Antimicrob Chemother* DOI:10.1093/jac/dkaa040.

32 Lee RS, Seemann T, Heffernan H, *et al.* Genomic epidemiology and antimicrobial resistance of Neisseria gonorrhoeae in New Zealand. *J Antimicrob Chemother* 2018; **73**: 353–64.

33 Mortimer TD, Pathela P, Crawley A, *et al.* The distribution and spread of susceptible and resistant Neisseria gonorrhoeae across demographic groups in a major metropolitan center. *Clin Infect Dis* DOI:10.1093/cid/ciaa1229.

34 Peng J-P, Yin Y-P, Chen S-C, *et al.* A Whole-genome Sequencing Analysis of Neisseria gonorrhoeae Isolates in China: An Observational Study. *EClinicalMedicine* 2019; **7**: 47–54.

35 Ryan L, Golparian D, Fennelly N, *et al.* Antimicrobial resistance and molecular epidemiology using whole-genome sequencing of Neisseria gonorrhoeae in Ireland, 2014–2016: focus on extended-spectrum cephalosporins and azithromycin. *Eur J Clin Microbiol Infect Dis* 2018; **37**: 1661–72.

36 Sánchez-Busó L, Golparian D, Corander J, *et al.* The impact of antimicrobials on gonococcal evolution. *Nature Microbiology* 2019; **4**: 1941–50.

37 Thomas JC, Seby S, Abrams AJ, *et al.* Evidence of Recent Genomic Evolution in Gonococcal Strains with Decreased Susceptibility to Cephalosporins or Azithromycin in the United States, 2014-2016. *J Infect Dis* DOI:10.1093/infdis/jiz079.

38 Town K, Harris S, Sánchez-Busó L, *et al.* Genomic and Phenotypic Variability in Neisseria gonorrhoeae Antimicrobial Susceptibility, England - Volume 26, Number 3—March 2020 - Emerging Infectious Diseases journal - CDC. DOI:10.3201/eid2603.190732.

39 Williamson DA, Chow EPF, Gorrie CL, *et al.* Bridging of Neisseria gonorrhoeae lineages across sexual networks in the HIV pre-exposure prophylaxis era. *Nat Commun* 2019; **10**: 1–10.

40 Yahara K, Nakayama S-I, Shimuta K, *et al.* Genomic surveillance of Neisseria gonorrhoeae to investigate the distribution and evolution of antimicrobial-resistance determinants and lineages. *Microb Genom* 2018; **4**. DOI:10.1099/mgen.0.000205.

41 Elwell LP, Roberts M, Mayer LW, Falkow S. Plasmid-Mediated Beta-Lactamase Production in Neisseria gonorrhoeae. *Antimicrobial Agents and Chemotherapy* 1977; published online March. DOI:10.1128/AAC.11.3.528.

42 Ropp PA, Hu M, Olesky M, Nicholas RA. Mutations in ponA, the Gene Encoding Penicillin-Binding Protein 1, and a Novel Locus, penC, Are Required for High-Level Chromosomally Mediated Penicillin Resistance in Neisseria gonorrhoeae. *Antimicrobial Agents and Chemotherapy* 2002; published online March. DOI:10.1128/AAC.46.3.769-777.2002.

43 Spratt BG, Bowler LD, Zhang Q-Y, Zhou J, Smith JM. Role of interspecies transfer of chromosomal genes in the evolution of penicillin resistance in pathogenic and commensal Neisseria species. *J Mol Evol* 1992; **34**: 115–25.

44 Dowson CG, Jephcott AE, Gough KR, Spratt BG. Penicillin-binding protein 2 genes of non-β-lactamase-producing, penicillin-resistant strains of Neisseria gonorrhoeae. *Molecular Microbiology* 1989; **3**: 35–41.

45 Brannigan JA, Tirodimos IA, Zhang Q-Y, Dowson CG, Spratt BG. Insertion of an extra amino acid is the main cause of the low affinity of penicillin-binding protein 2 in penicillin-resistant strains of Neisseria gonorrhoeae. *Molecular Microbiology* 1990; **4**: 913–9.

46 Bharat A, Demczuk W, Martin I, Mulvey MR. Effect of Variants of Penicillin-Binding Protein 2 on Cephalosporin and Carbapenem Susceptibilities in Neisseria gonorrhoeae. *Antimicrobial Agents and Chemotherapy* 2015; published online May 18. https://journals.asm.org/doi/abs/10.1128/AAC.05143-14 (accessed Dec 17, 2021).

47 Tomberg J, Unemo M, Davies C, Nicholas RA. Molecular and Structural Analysis of Mosaic Variants of Penicillin-Binding Protein 2 Conferring Decreased Susceptibility to Expanded-Spectrum Cephalosporins in Neisseria gonorrhoeae: Role of Epistatic Mutations. *Biochemistry* 2010; **49**: 8062–70.

48 Morse SA, Johnson SR, Biddle JW, Roberts MC. High-level tetracycline resistance in Neisseria gonorrhoeae is result of acquisition of streptococcal tetM determinant. *Antimicrobial Agents and Chemotherapy* 1986; published online Nov. DOI:10.1128/AAC.30.5.664.

49 Hu M, Nandi S, Davies C, Nicholas RA. High-Level Chromosomally Mediated Tetracycline Resistance in Neisseria gonorrhoeae Results from a Point Mutation in the rpsJ Gene Encoding Ribosomal Protein S10 in Combination with the mtrR and penB Resistance Determinants. *Antimicrobial Agents and Chemotherapy* 2005; published online Oct. DOI:10.1128/AAC.49.10.4327-4334.2005.

50 Sparling PF, Sarubbi FA, Blackman E. Inheritance of low-level resistance to penicillin, tetracycline, and chloramphenicol in Neisseria gonorrhoeae. *J Bacteriol* 1975; **124**: 740–9.

51 Gill MJ, Simjee S, Al-Hattawi K, Robertson BD, Easmon CSF, Ison CA. Gonococcal Resistance to β-Lactams and Tetracycline Involves Mutation in Loop 3 of the Porin Encoded at thepenB Locus. *Antimicrob Agents Chemother* 1998; **42**: 2799–803.

52 Xia M, Whittington WLH, Shafer WM, Holmes KK. Gonorrhea among Men Who Have Sex with Men: Outbreak Caused by a Single Genotype of Erythromycin-Resistant Neissevia gonorrhoeae with a Single-Base Pair Deletion in the mtrR Promoter Region. *J Infect Dis* 2000; **181**: 2080–2.

53 Veal WL, Nicholas RA, Shafer WM. Overexpression of the MtrC-MtrD-MtrE Efflux Pump Due to an mtrR Mutation Is Required for Chromosomally Mediated Penicillin Resistance in Neisseria gonorrhoeae. *Journal of Bacteriology* 2002; **184**: 5619–24.
